# Ciliary integrity defines a central canal repair checkpoint linking microtubule stabilization to spinal cord regeneration

**DOI:** 10.64898/2026.06.17.733052

**Authors:** Chunnuan Lin, Rui Zhang, Xianming Wu, Lamei Yang, Honglin Xu, Ruifan Lin, Yannan Zhao, Qi Xie, Jianwu Dai, Wenxiang Meng

**Affiliations:** Laboratory of Integrative Physiology, Institute of Genetics and Developmental Biology, Chinese Academy of Sciences, Beijing 100101, China; University of Chinese Academy of Sciences, Beijing 100049, China; State Key Laboratory of Female Fertility Promotion, Department of Traditional Chinese Medicine, Peking University Third Hospital, Beijing 100191, China; Wangjing Hospital of China Academy of Chinese Medical Sciences, Beijing 100102, China; Tianjin Key Laboratory of Biomedical Materials, Institute of Biomedical Engineering, Chinese Academy of Medical Sciences and Peking Union Medical College, Tianjin 300192, China

## Abstract

Microtubule-stabilizing agents consistently improve functional recovery after spinal cord injury (SCI), yet the structural mechanism underlying their shared therapeutic effects remains unclear. Here, we find that chemically distinct stabilizers converge on preservation of ciliary integrity within central canal-associated cells, including ependymal cells and cerebrospinal fluid–contacting neurons. Using complementary SCI models, including complete transection and crush injury, we observe that maintenance of ciliary architecture is associated with reduced glial scarring, improved tissue continuity, and enhanced locomotor recovery. Single-cell transcriptomic analysis further identifies these cell populations as prominent responders to microtubule stabilization, with ciliogenesis-related programs selectively preserved. Importantly, pharmacological disruption of cilia-associated signaling attenuates recovery, whereas promoting ciliogenesis partially recapitulates therapeutic effects, identifying ciliary integrity as a critical cilia-associated structural dependency that contributes to microtubule-stabilizer-mediated spinal cord repair. Together, these findings identify a cilia-dependent central canal regenerative niche as a candidate structural checkpoint linking microtubule stabilization to functional recovery after SCI and identify ciliogenesis as a therapeutically actionable target for SCI.

**Graphical abstract:** 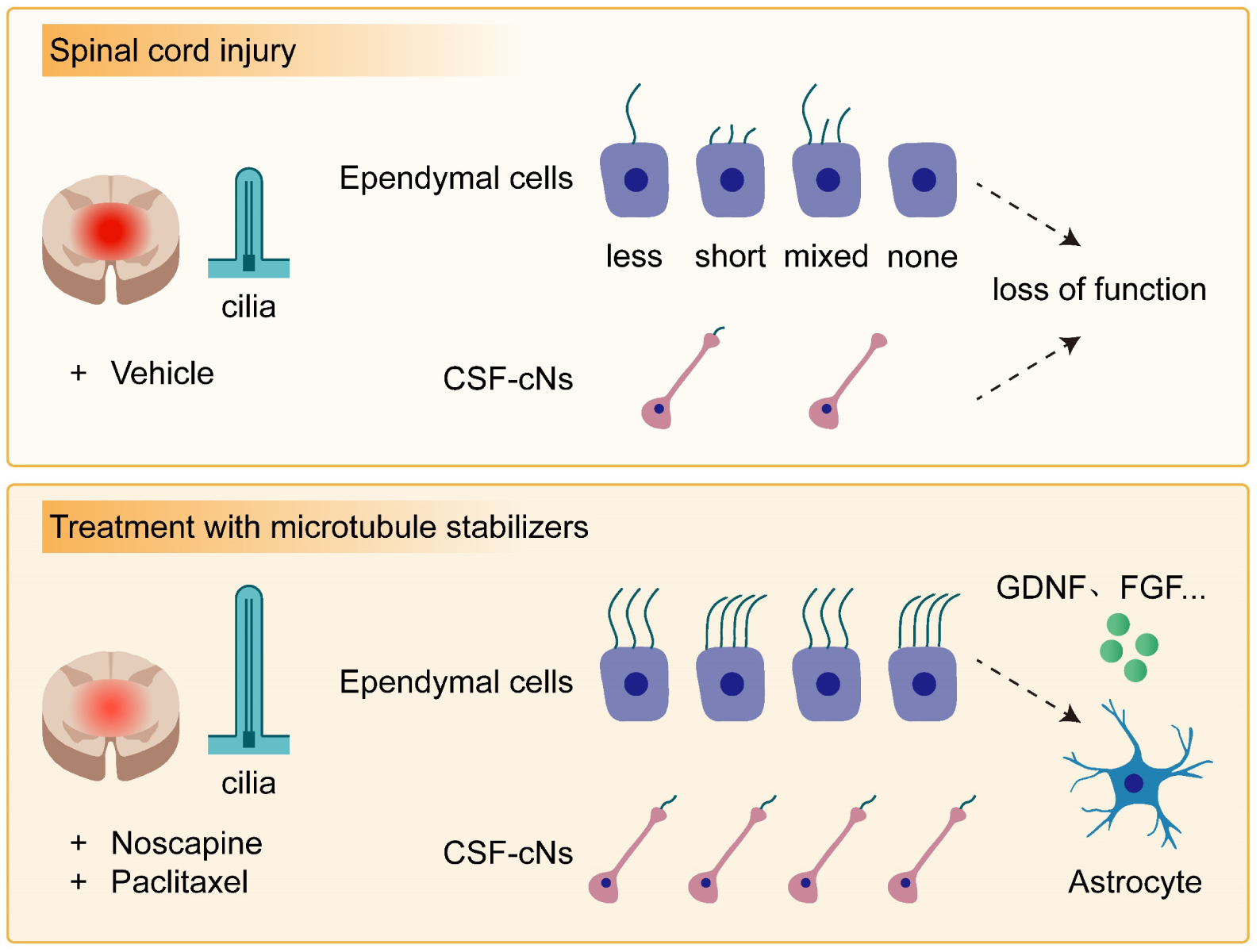

## Introduction

Spinal cord injury (SCI) remains one of the most formidable challenges in regenerative neuroscience, as the primary trauma is rapidly followed by secondary pathological cascades-including inflammation, vascular dysfunction, cytoskeletal collapse, and glial barrier formation-that severely restrict structural and functional recovery^1,2^. Although several experimental interventions have shown promise, strategies capable of simultaneously rebuilding structure and restoring function remain limited^3–5^, underscoring the importance of identifying broadly targetable structural processes.

Microtubule-stabilizing agents have emerged as encouraging candidates for SCI repair^6,7^. Compounds such as Paclitaxel and Epothilones, which bind to the taxane pocket within the β-tubulin lumen^7,8^, promote axon regeneration, reduce fibrotic scarring, and improve locomotor outcomes in preclinical models^9–11^. These results suggest that stabilizing the microtubule cytoskeleton may exert broad reparative effects; however, the mechanistic basis of their shared benefits remains unclear. Given the substantial differences in chemotype and binding mode among these stabilizers, it is unknown whether they act through unrelated pathways or converge on a common downstream structural process. This unresolved question not only limits our conceptual understanding of structural interventions but also constrains the rational development of effective therapies.

To address this gap, we systematically compared microtubule stabilizers with divergent structures in a complete transection model and found that all improved functional recovery to varying degrees, suggesting the presence of a structure-independent downstream mechanism. We then selected Noscapine-a non-taxane stabilizer^12,13^-for in-depth mechanistic dissection, integrating Brainbow-based circuit tracing, histology, behavioral assessment, and single-cell RNA sequencing. Unbiased transcriptomic profiling identified two central canal-associated populations-ependymal cells and cerebrospinal fluid-contacting neurons (CSF-cNs)-as the principal responders to microtubule stabilization.

Transcriptomic signatures in both populations consistently emphasized cilia-related programs, a conceptually compelling finding given that cilia are microtubule-based organelles essential for cerebrospinal fluid flow, environmental sensing, and lineage regulation^14,15^. These observations led us to hypothesize that cilia integrity may serve as a structural bridge linking microtubule stabilization to cell-type-specific reparative responses. In support of this, Noscapine preserved ciliary structure in both cell types in vivo, while pharmacological disruption of cilia-dependent signaling markedly reduced its cellular and functional benefits. Here, we tested the hypothesis that microtubule stabilization promotes spinal cord repair by maintaining a cilia-defined central canal regenerative niche that acts as a structural checkpoint linking cytoskeletal stability to functional recovery after injury.

Together, these findings identify the preservation of ciliary integrity as a shared structural checkpoint underlying microtubule-based spinal cord repair. By connecting microtubule stabilization to the regenerative functions of central canal-associated cells, our work defines a cilia-dependent central canal niche as a mechanistic axis of repair and emphasizes ciliogenesis as a practical and broadly applicable therapeutic target for SCI.

## Results

### Structurally distinct microtubule stabilizers converge on a shared structural mechanism of tissue repair and locomotor recovery

We compared five microtubule stabilizers spanning distinct chemotypes and binding pockets-including Noscapine (non-taxane, colchicine-related^12,13^), Paclitaxel and Epothilone D (taxane pocket^16,17^), TPI-287 (taxane derivative^18^), and TTI-237 (vinca-like binder^19^) -in a complete transection spinal cord injury (SCI) model (Figures 1A and 1B). All compounds reduced EB1⁺ comet density in vivo (Figures S1A and S1B), confirming suppression of microtubule dynamics as a shared upstream action. Despite their divergent structures and binding modes, each stabilizer improved locomotor recovery at 4 weeks post-injury (wpi) (Figure 1C). These cross-chemotype benefits suggest that stabilizers acting through diverse microtubule-binding sites may converge on a common downstream process.

**Figure 1.**
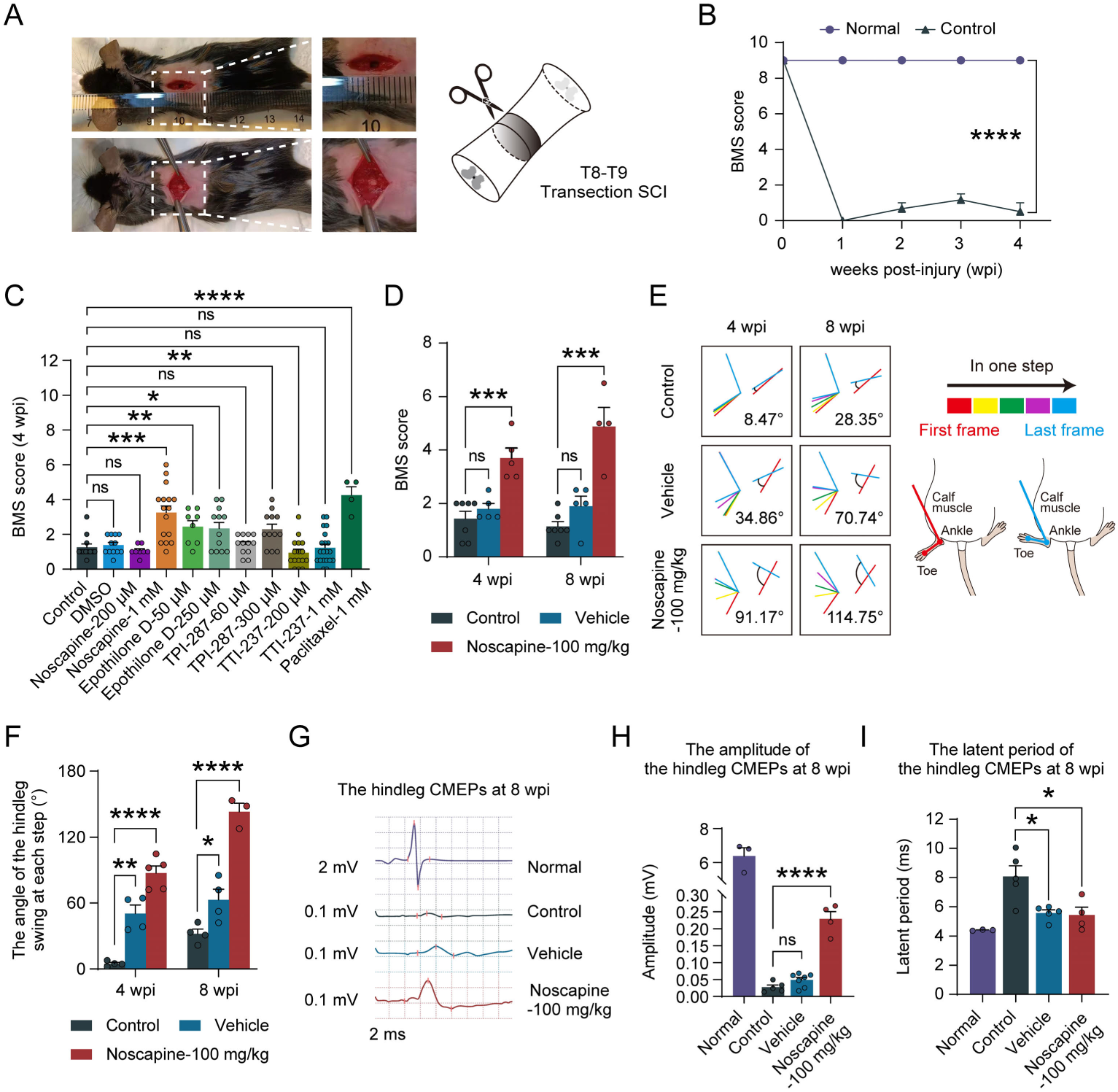
Distinct chemotypes of microtubule stabilizers promote functional recovery in a complete transection model. **(A)** The diagram of surgical procedures of thoracic complete transection spinal cord injury. **(B)** The BMS score of the hindleg of normal mice and control group mice within 4 weeks post-injury (wpi) (n = 3 per group). **(C)** The BMS score of the hindleg of mice treated with microtubule stabilizers combined with a linear-ordered collagen scaffold (LOCS) at 4 wpi (n = 12 for Control, n = 12 for DMSO, n = 8 for Noscapine-200 μM, n = 16 for Noscapine-1 mM, n = 8 for Epothilone D-50 μM, n = 12 for Epothilone D-250 μM, n = 12 for TPI-287-60 μM, n = 12 for TPI-287-300 μM, n = 16 for TTI-237-200 μM, n = 20 for TTI-237-1 mM, n = 4 for Paclitaxel-1 mM). **(D)** The BMS score at 4 and 8 wpi with intraperitoneal injection (4 wpi, n = 7 for Control, n = 5 for Vehicle and Noscapine-100 mg/kg; 8 wpi, n = 7 for Control, n = 5 for Vehicle, n = 4 for Noscapine-100 mg/kg). **(E)** The representative images of simulated mouse hindleg swing at 4 and 8 wpi. The line from the toe to the calf muscle of the lower leg is used to represent the anterior-posterior swing of the hindleg when taking one step. **(F)** Quantitative analysis of the angle of the hingleg swing at each step at 4 and 8 wpi (4 wpi, n = 4 for Control and Vehicle, n = 5 for Noscapine-100 mg/kg; 8 wpi, n = 4 for Control and Vehicle, n = 3 for Noscapine-100 mg/kg). **(G)** The representative images of cortical motor evoked potentials (CMEPs) between normal mice and experimental group mice at 8 wpi. **(H-I)** Quantitative analysis of the amplitude **(H)** (n = 3 for Normal, n = 5 for Control, n = 7 for Vehicle, n = 4 for Noscapine-100 mg/kg) and latent period **(I)** (n = 3 for Normal, n = 5 for Control, n = 5 for Vehicle, n = 4 for Noscapine-100 mg/kg) of the hindleg CMEPs at 8 wpi. SCI, spinal cord injury, wpi, weeks post-injury. Data are shown as mean ± SEM. ns, not significant (*P* > 0.05), \**P* < 0.05, \*\**P* < 0.01, \*\*\**P* < 0.001, \*\*\*\**P* < 0.0001. Unpaired two-sided Student’s t test.

Given that Noscapine is a non-taxane microtubule stabilizer with a favorable safety profile and reported CNS accessibility, and produced robust motor recovery in our screening, this study selected Noscapine as a typical microtubule stabilizer and further screened its optimal concentration for intraperitoneal injection. Dose-response testing identified 100 mg/kg Noscapine as the optimal regimen, yielding the strongest behavioral and cortical motor evoked potentials (CMEPs) improvements at 4 wpi (Figures S1C-S1F). Assessments of motor function (Figure 1D), hindlimb range of motion (Figures 1E-F) and electrophysiological examinations (Figures 1G-I) performed at 8 wpi demonstrated that the therapeutic effects of Noscapine were sustained and stable.

To evaluate how microtubule stabilization influences tissue repair, we used a unified paradigm in a thoracic complete transection SCI model involving local implantation of a linear-ordered collagen scaffold (LOCS) and systemic drug administration (Figure 2A). Laminin staining at 4 wpi showed reduced ECM/fibrotic scarring in both Noscapine- and Paclitaxel-treated cords (Figures 2B-2D). NeuN and SMI-312 labeling revealed increased neuronal presence and more continuous nerve fibers across the lesion in both Noscapine and Paclitaxel groups (Figures 2E-2H).

**Figure 2.**
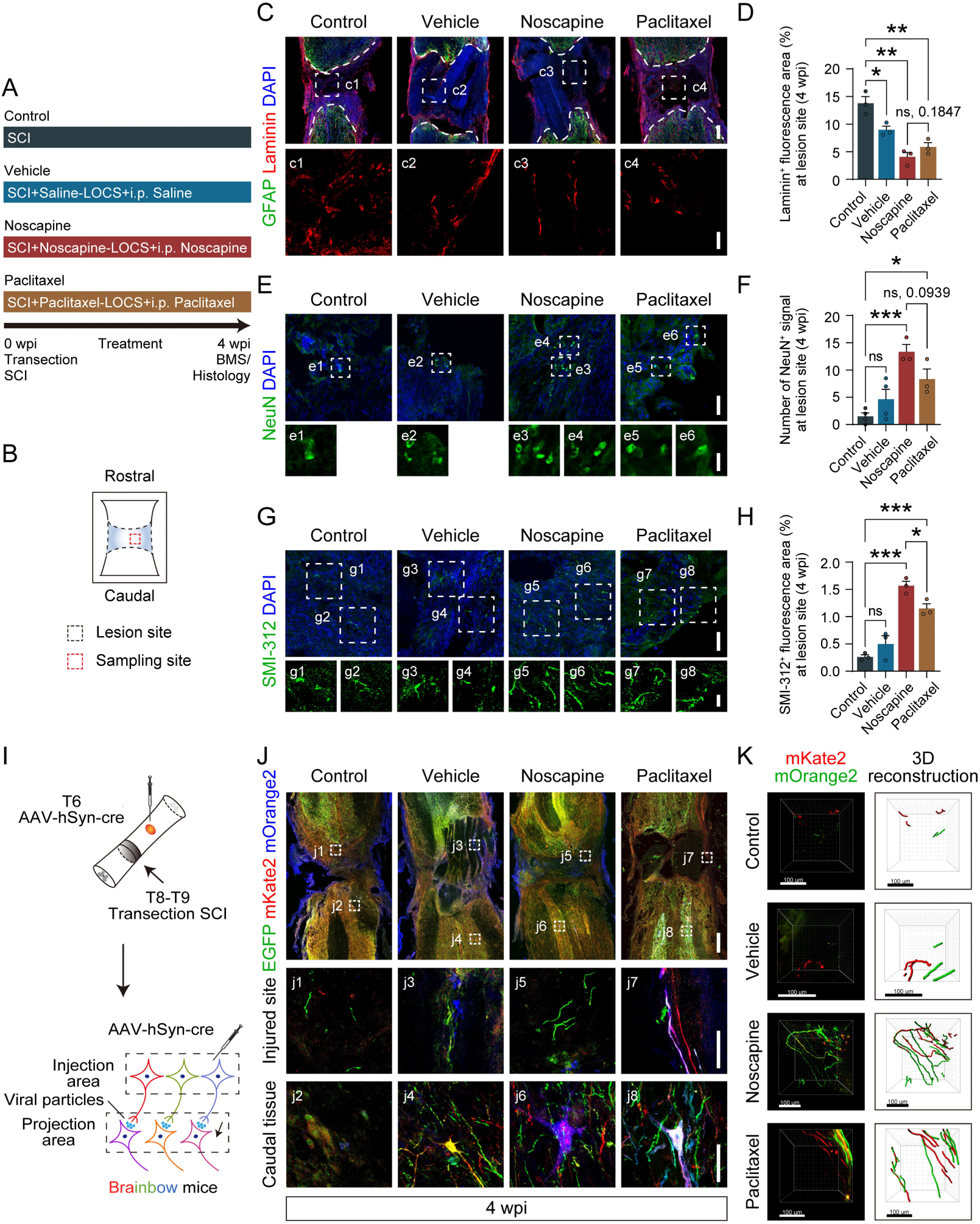
Microtubule stabilizers attenuate scarring and enhance anatomical continuity across the lesion. **(A)** Diagram of experimental grouping and operation timeline. **(B)** The schematic diagram of the location of the immunostaining images in C, E, and G. **(C)** Immunostaining of GFAP and Laminin showing the ECM/fibrotic scar in the lesion site at 4 wpi, the details of Laminin in the white squares are shown in the enlarged images (c1, c2, c3, c4). Scale bars, 200 μm on the top and 50 μm on the bottom. **(D)** Quantitative analysis of the proportion of Laminin^+^ fluorescence area in the lesion site at 4 wpi for each group (n = 3 per group). **(E)** Immunostaining of NeuN showing the survival neurons in the lesion site at 4 wpi, the details of NeuN in the white squares are shown in the enlarged images (e1, e2, e3, e4, e5, d6). Scale bars, 50 μm on the top and 20 μm on the bottom. **(F)** Quantitative analysis of the number of NeuN^+^ signal in the lesion site at 4 wpi for each group (n = 4 for Control and Vehicle, n = 3 for Noscapine and Paclitaxel). **(G)** Immunostaining of SMI-312 showing the nerve fibers in the lesion site at 4 wpi, the details of SMI-312 in the white squares are shown in the enlarged images (g1, g2, g3, g4, g5, g6, g7, g8). Scale bars, 50 μm on the top and 20 μm on the bottom. **(H)** Quantitative analysis of the proportion of SMI-312^+^ fluorescence area in the lesion site at 4 wpi for each group (n = 3 per group). **(I)** Diagram of random multicolor marking scheme based on Brainbow system and anterograde trans-synaptic AAV virus. **(J)** Immunostaining of EGFP, mKate2 and mOrange2 showing the tracing results using Brainbow system at 4 wpi, the labeling details of injured site were shown in j1, j3, j5, j7, the caudal tissue were shown in j2, j4, j6, j8. Scale bars, 300 μm on the top and 50 μm on the bottom. **(K)** Immunostaining of mKate2 and mOrange2 and tissue clearing showing the 3D rendering and reconstructed images of continuous neurons in the lesion site at 4 wpi. Scale bars, 100 μm. Staining of EGFP was not shown because the signal was lost during tissue clearing. SCI, spinal cord injury, i.p., intraperitoneal injection. Data are shown as mean ± SEM. ns, not significant (*P* > 0.05), \**P* < 0.05, \*\**P* < 0.01, \*\*\**P* < 0.001. Unpaired two-sided Student’s t test.

To determine whether axon-like fibers extended across the lesion, we performed anterograde labeling in Brainbow mice using AAV-hSyn-Cre^20,21^ (Figures 2I and S2). Both stabilizers supported multicolor axon-like fibers extending across the lesion and neuronal somata caudal to the injury (Figure 2J). Whole-cord clearing further revealed tract-like structures that appeared to span the lesion, providing structural substrates for cross-lesion axonal continuity (Figure 2K).

These findings show that microtubule stabilizers with distinct structures could promote the recovery of motor function, attenuate fibrotic scarring and support structural substrates for cross-lesion circuit continuity. Their consistent effects across distinct chemotypes extend previous taxane-based observations and suggest that microtubule stabilization plays an essential role in the therapeutic intervention for SCI.

### Noscapine modulates the early lesion microenvironment after SCI, as revealed by single-cell transcriptomics

To determine whether Noscapine influences early cellular states after SCI, we performed whole-cell single-cell RNA sequencing (scRNA-seq) on lesion-core tissue collected at 2 and 3 wpi from thoracic complete transection models (Figures 3A, S3 A, and S3B). Following quality control, 51,117 cells were retained (Figures S3C-S3E), resolving 14 major cell types and five proliferative subclusters (Figures S4A-S4C) spanning immune, vascular/stromal, and neurocyte compartments (Figures S4D, S5A, and 3B).

**Figure 3.**
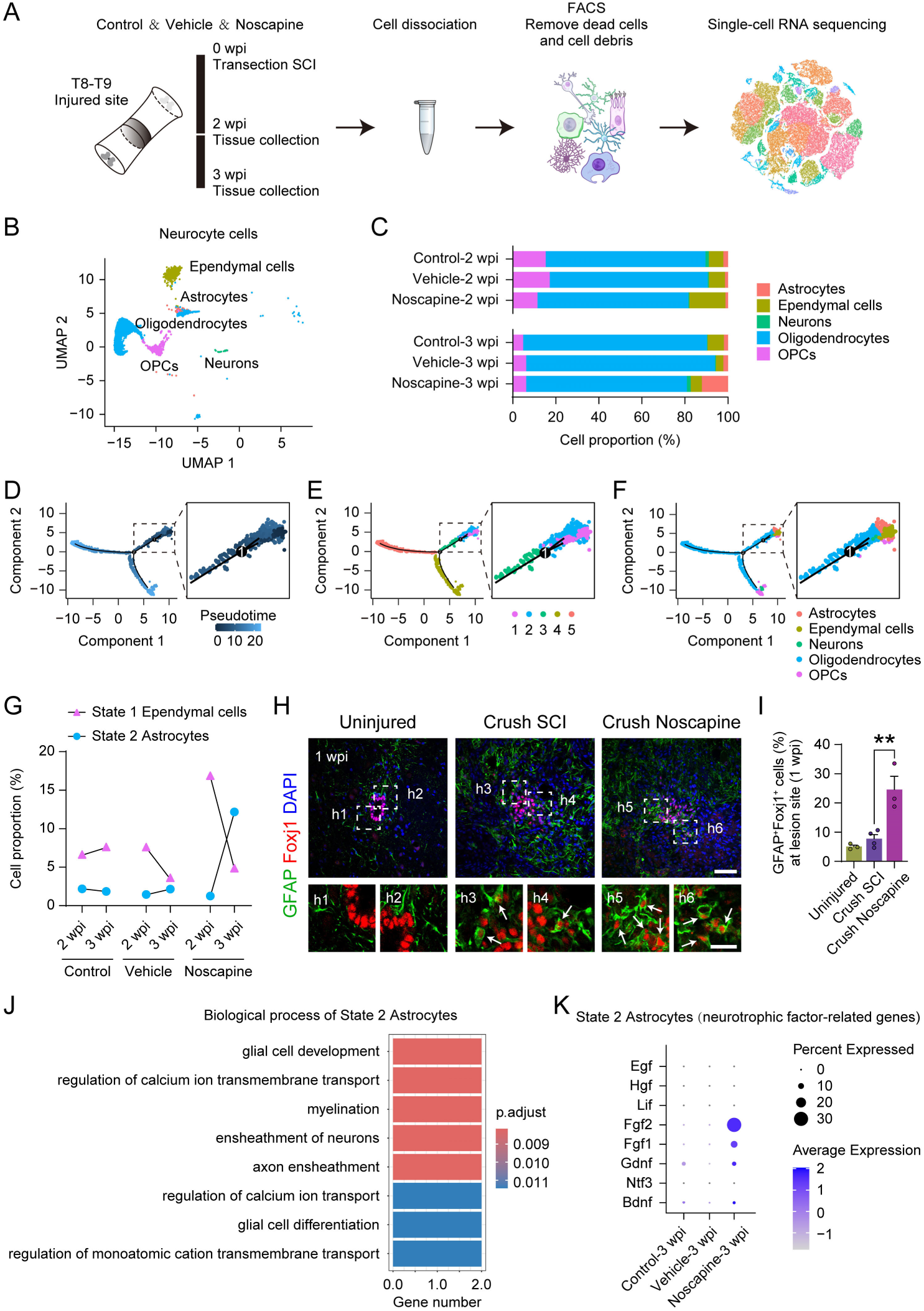
Noscapine activates the ependymal cells and induces differentiation towards a functional state. **(A)** Overview of the experimental approach for single-cell RNA sequencing analysis of the spinal cord lesion site tissue from three experimental groups at 2 wpi (subacute phase) and 3 wpi (chronic phase). **(B)** UMAP plot showing neurocyte cells (astrocytes, ependymal cells, neurons, oligodendrocytes, OPCs) of six groups. **(C)** The percentage of neurocyte cells in each group. **(D-F)** Single-cell trajectory of neurocyte cells using Monocle, it separately demonstrates the pseudotime of cells **(D)**, the state of cells on the trajectory tree **(E)**, and the distribution characteristics of the five cell types **(F)**. **(G)** The percentage of State 1 Ependymal cells and State 2 Astrocytes among the groups. **(H)** Immunostaining of GFAP and Foxj1 showing the ependymal cells exhibiting GFAP-labeled characteristics in the lesion site at 1 week post crush injury, the details of the double-positive cells in the white squares are shown in the enlarged images (h1, h2, h3, h4, h5, h6). Scale bars, 50 μm on the top and 20 μm on the bottom. **(I)** Quantitative analysis of the proportion of GFAP^+^Foxj1^+^ cells among Foxj1^+^ cells in the lesion site at 1 wpi for each group (n = 3 for Uninjured and Crush Noscapine, n = 4 for Crush SCI). **(J)** GO enrichment analysis of biological process for the differentially expressed genes in State 2 Astrocytes compared to State 1 Astrocytes. **(K)** Dot plot showing the expression of neurotrophic factor-related genes in State 2 Astrocytes across each group at 3 wpi. The dot size indicates the percentage of cells in which that gene is detected, and the color represents the average expression level. Data are shown as mean ± SEM. \*\**P* < 0.01. Unpaired two-sided Student’s t test.

Noscapine reshaped both immune and vascular/stromal niches. Microglia, which constituted >60% of cells in control, declined to < 5% at 3 wpi in Noscapine-treated samples (Figures S4D and S4E), Although changes in dissociation sensitivity and lesion composition cannot be fully excluded, the reduced representation of inflammatory populations was consistent with histological evidence of attenuated lesion barriers^22–24^. Fibroblast populations also decreased (Figures S5A and S5B), particularly extracellular matrix-enriched clusters (clusters 1-2; Figures S5C-S5F). Angiogenic (cluster 0) and proliferative (cluster 3) fibroblast subsets shifted as well (Figures S5G and S5H), aligning with the reduction of laminin deposition observed histologically (Figures 2C and 2D).

Thus, Noscapine rapidly alleviates inflammatory and fibrotic constraints early after injury, establishing a more permissive microenvironment for repair. However, microenvironmental modulation alone cannot explain the convergent actions of structurally distinct microtubule stabilizers, highlighting the need to identify downstream pathways directly tied to microtubule stabilization.

### Noscapine activates central canal**-**associated cell populations and promotes their reparative functions

SCI induces pronounced shifts in neurocyte composition: neurons, astrocytes, and oligodendrocytes are depleted, whereas previously quiescent populations become activated and recruited to the lesion site^25,26^. We focused on five neurocyte classes-astrocytes, ependymal cells, neurons, oligodendrocytes, and oligodendrocyte precursor cells (OPCs) (Figure 3B). Between 2 and 3 wpi, Noscapine-treated cords exhibited an increase in astrocytes accompanied by a decrease in ependymal cells, a coordinated pattern not observed in controls (Figure 3C). Given the stem-like nature of adult ependymal cells^27–29^, these complementary changes suggested altered lineage dynamics.

Pseudotime reconstruction revealed two major trajectories from ependymal cells-toward astrocytes (State 2) and oligodendrocytes (State 3)-with undifferentiated cells corresponding to State 1 (Figures 3D-3F). Noscapine reduced State 1 and expanded State 2 over time (Figure 3G). No similar shifts were observed in other states (Figure S6A). Because complete transection disrupts central canal organization, we validated the differentiation trend using a thoracic crush SCI model (Figures S6B and S6C)^30,31^. At 1 wpi, the proportion of GFAP/Foxj1 double-positive cells were significantly increased following Noscapine treatment (Figures 3H and 3I). Specifically, the percentage of ependymal cell-associated astrocytes increased significantly. These results suggest that Noscapine intervention modulates the function of ependymal cells, and is associated with an increased emergence of ependymal cell-associated astrocytes (GFAP^+^Foxj1^+^ cells) in the lesion core after SCI, although formal demonstration of an ependymal-to-astrocyte lineage transition will require dedicated lineage tracing.

State 2 astrocytes were enriched for processes related to glial development and neuronal ensheathment (Figure 3J). By the chronic phase, Noscapine-treated astrocytes expressed higher levels of neurotrophic genes-including Bdnf, Gdnf, Fgf1, and Fgf2 (Figure 3K)-indicating a reparative rather than scar-forming phenotype^32,33^.

To characterize neuron subtypes relevant to functional recovery after spinal cord injury, we compared our preliminarily identified neurons with the annotated thoracic neuron profiles reported by Squair J.W. et al. using single-nucleus sequencing^34^. Reclustering of neuronal subsets identified cerebrospinal fluid-contacting neurons (CSF-cNs) as PKD2L1-expressing specialized neurons located around the central canal (Figures S7A and S7B)^14^. At 4 weeks after complete transection SCI, CSF-cNs were detected in the noscapine group (Figure S7C), indicating that CSF-cNs may serve as a key responsive neuronal subtype contributing to functional recovery following treatment with Noscapine. We further performed immunostaining for PKD2L1 at 1 week post crush SCI, and the statistical results verified that Noscapine enhanced the survival of CSF-cNs (Figures S7D and S7E).

Together, these findings show that Noscapine exerts dual actions on central canal-associated cells-promoting the emergence of ependymal cell-associated, repair-associated astrocytes and preserving CSF-cNs. These coordinated responses reshape the canal microenvironment and provide a cellular basis for improved tissue remodeling and repair after SCI.

### Microtubule stabilizers preserve ciliary integrity of central canal**-**associated cells after SCI

To investigate the basis of Noscapine’s effects on central canal-associated cells, we examined the transcriptional changes of these cells. Gene ontology enrichment revealed that State 1 ependymal cells were strongly enriched for cilia-related biological processes-including cilium movement, cilium- or flagellum-dependent motility, and cilium assembly (Figure 4A). These pathways are central to the structural and sensory roles of ependymal cilia in cerebrospinal fluid dynamics and lineage activation^35,36^. At the gene level, components essential for ciliary assembly (Arl13b, Ccdc114) and intraflagellar transport (Ift122, Ift46) were upregulated in Noscapine-treated samples, particularly at 2 wpi (Figure 4B). Because Noscapine is not known to act as a transcriptional activator, these transcriptional signatures likely reflect preserved cellular structure that maintains the capacity for cilia-associated pathways. Ependymal cells line the central canal and extend motile multicilia into cerebrospinal fluid^36,37^. After SCI, these cilia commonly undergo fragmentation and shedding, disrupting both organization and sensory function^38^. To directly test whether Noscapine protects ciliary structure, we performed immunostaining in the thoracic crush model, where central canal architecture is preserved as much as possible. At 1 wpi, Noscapine-treated cords displayed a significantly higher proportion of ciliated ependymal cells and showed continuous, organized ciliary arrays rather than fragmented clusters (Figures 4C and 4D), indicating early protection against injury-induced cilia deterioration.

**Figure 4.**
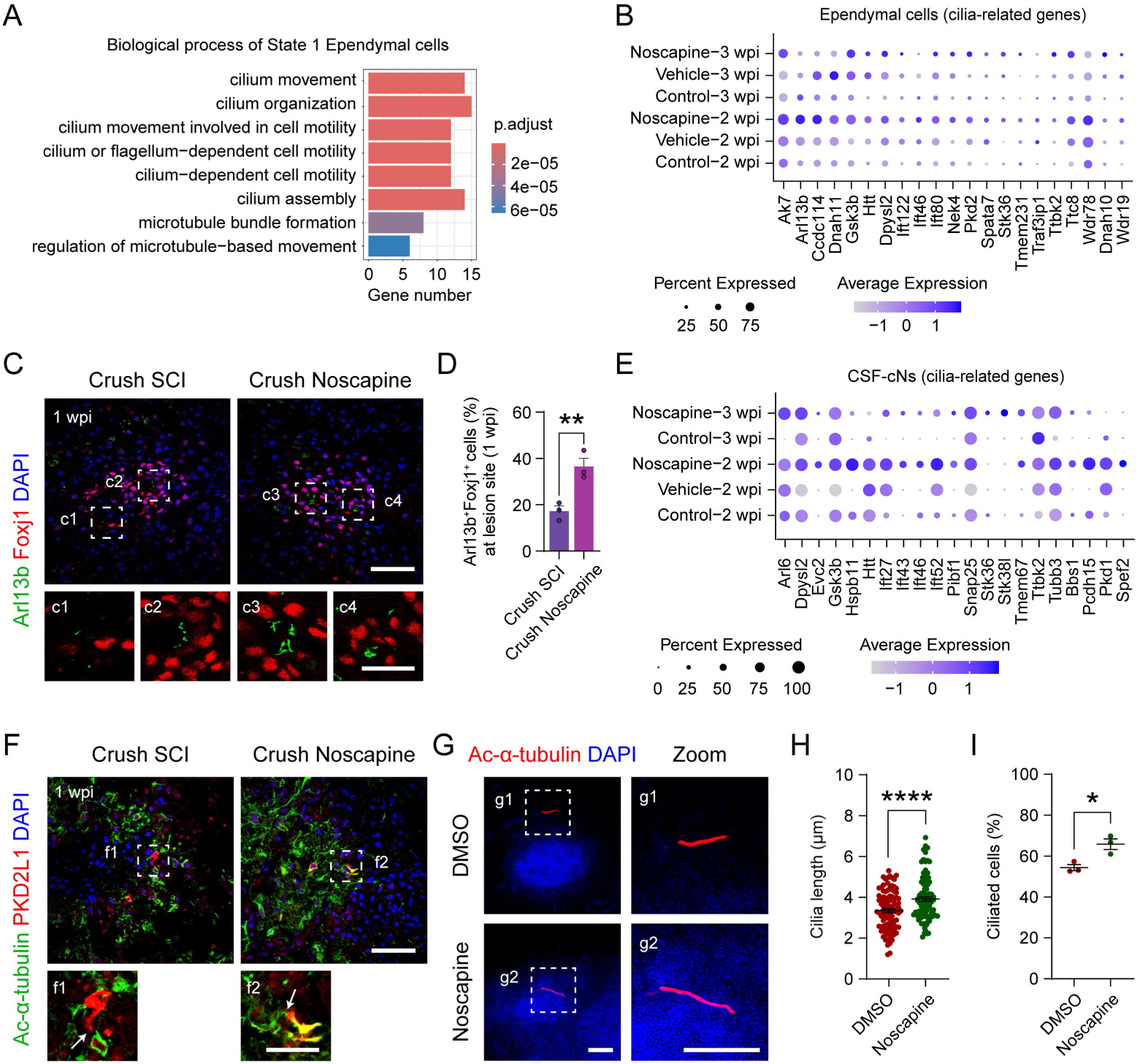
Noscapine preserves ciliary integrity of central canal–associated cells after SCI. **(A)** GO enrichment analysis of biological process for the differentially expressed genes in State 1 Ependymal cells compared to State 2 Ependymal cells. **(B)** Dot plot showing the expression of cilia-related genes in ependymal cells across each group. The dot size indicates the percentage of cells in which that gene is detected, and the color represents the average expression level. **(C)** Immunostaining of Arl13b and Foxj1 showing the ependymal cells with ciliary features in the lesion site at 1 wpi, the details of the double-positive cells in the white squares are shown in the enlarged images (c1, c2, c3, c4). Scale bars, 50 μm on the top and 20 μm on the bottom. **(D)** Quantitative analysis of the proportion of Arl13b^+^Foxj1^+^ cells among Foxj1^+^ cells in the lesion site at 1 wpi for each group (n = 3 per group). **(E)** Dot plot showing the expression of cilia-related genes in CSF-cNs across each group. The dot size indicates the percentage of cells in which that gene is detected, and the color represents the average expression level. **(F)** Immunostaining of acetylated-tubulin (Ac-α-tubulin) and PKD2L1 showing the CSF-cNs with ciliary features in the lesion site at 1 wpi, the details of the double-positive cells in the white squares are shown in the enlarged images (f1, f2). Scale bars, 50 μm on the top and 20 μm on the bottom. **(G)** Immunostaining of Ac-α-tubulin showing the effect of Noscapine on the cilia in hTERT-RPE1 cells. Scale bar, 5 μm. **(H)** Quantitative analysis of the length of cilia in each treatment (n = 100 per group). **(I)** Quantitative analysis of the percentage of ciliated cells in each treatment (n = 3 per group). Data are shown as mean ± SEM. \**P* < 0.05, \*\**P* < 0.01, \*\*\*\**P* < 0.0001. Unpaired two-sided Student’s t test.

CSF-cNs represent another ciliated cell type surrounding the central canal^14,39^. These PKD2L1-positive neurons sense cerebrospinal fluid composition and modulate locomotor circuits^40^, and they are known to be highly vulnerable after SCI^41,42^. Noscapine increased expression of cilia-associated genes such as Ift43 and Ift46 within CSF-cNs (Figure 4E), and immunostaining confirmed that a greater proportion of CSF-cNs retained intact cilia following treatment (Figure 4F).

To assess whether Noscapine can directly influence ciliary structure outside the injury environment, we analyzed human retinal pigment epithelial (hTERT RPE-1) cells. Within 24 hours of treatment, Noscapine increased cilia length and the proportion of ciliated cells (Figures 4G-4I). Because RPE-1 cells are not subject to injury-induced ciliary fragmentation, these findings suggest a direct capacity of Noscapine to support ciliary structure in vitro. Altogether, Noscapine protects ciliary structure and function in both ependymal cells and CSF-cNs after SCI. Preservation of these microtubule-based organelles likely maintains cerebrospinal fluid flow, environmental sensing, and lineage competence, thereby supporting structural and functional recovery.

Paclitaxel, a widely acknowledged taxane microtubule stabilizer, exhibits distinct differences from Noscapine in chemical structure and tubulin binding sites; nevertheless, the two compounds share similar microtubule stabilizing activities^43,44^. To further explore whether Paclitaxel shares a similar mechanism with Noscapine, we treated mice with intraperitoneally injected Paclitaxel in the crush SCI model (Figure S8A). Statistical analyses revealed that at 1 wpi, the survival of CSF-cNs, the proportion of ependymal cell-associated astrocytes, as well as the percentage of ependymal cells with ciliary structures were all significantly increased (Figure S8B–G). Therefore, reinforcing ciliary integrity serves as a cross-chemotype structural effector of microtubule-based repair.

These findings establish ciliary integrity as a downstream structural effector of microtubule stabilization and suggest that it may serve as a unifying mediator of microtubule stabilizers’ reparative actions.

### Cilia represent a critical structural intervention target for facilitating spinal cord injury repair

To directly determine whether the reparative effects of Noscapine depend on ciliary function, we used SANT-1, a Smoothened receptor antagonist that disrupts Sonic hedgehog (Shh) signaling and downstream cilia-dependent regulation^45^, thereby mimicking ciliary impairment (Figures 5A-5C). In the thoracic crush model, co-administration of SANT-1 partially reversed the locomotor improvements induced by Noscapine alone (Figures 5D and 5E), indicating that Noscapine’s functional benefits are at least partly cilia-dependent. At the tissue level, SANT-1 markedly reduced Arl13b-positive cilia in ependymal cells (Figures 5F and 5G), confirming effective disruption of ciliary maintenance.

**Figure 5.**
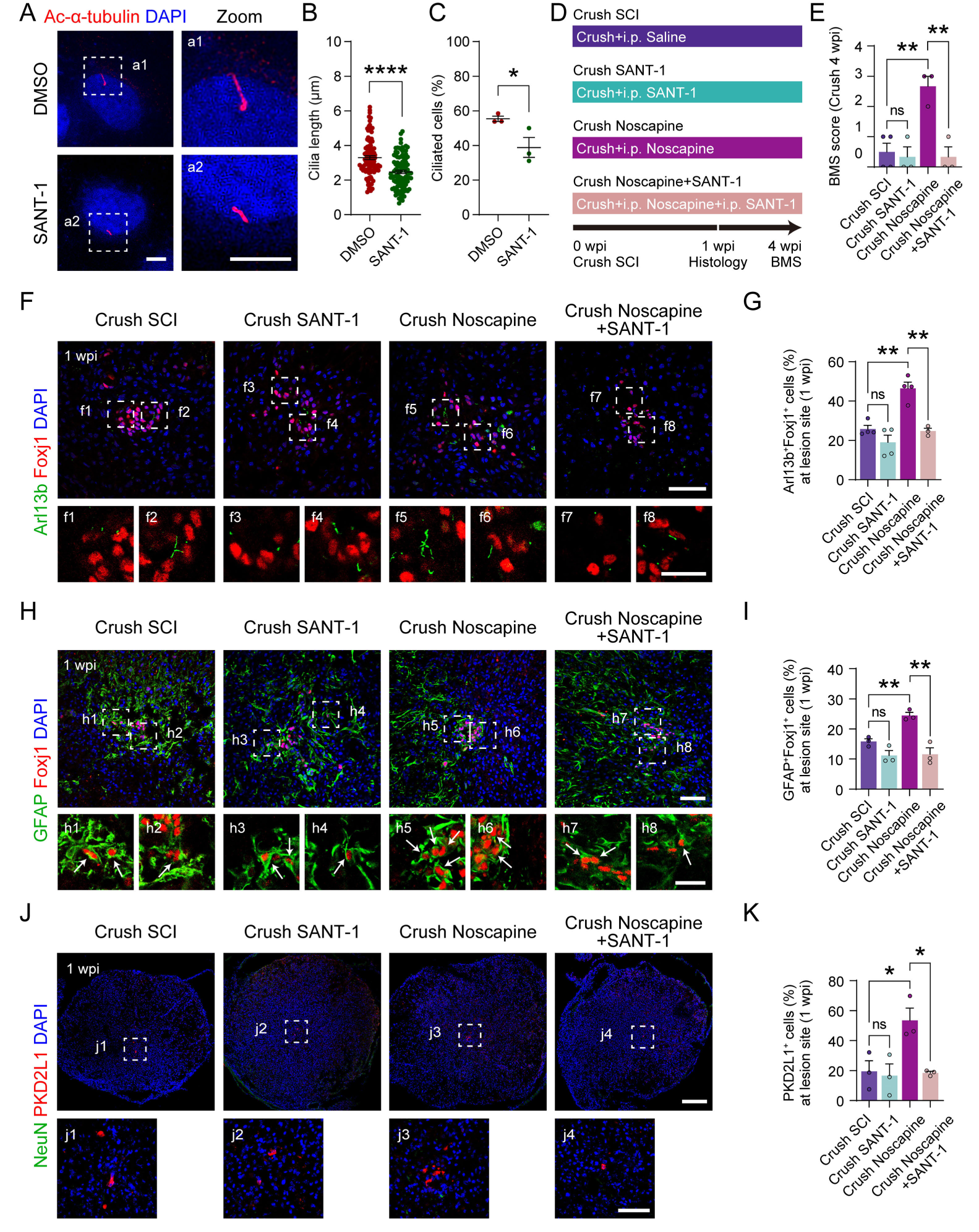
Ciliary integrity is required for the functional recovery mediated by Noscapine. **(A)** Immunostaining of acetylated-tubulin (Ac-α-tubulin) showing the effect of SANT-1 on the cilia in hTERT-RPE1 cells. Scale bar, 5 μm. (B) Quantitative analysis of the length of cilia in each treatment (n = 100 per group). **(C)** Quantitative analysis of the percentage of ciliated cells in each treatment (n = 3 per group). **(D)** Diagram of experimental grouping and operation timeline using SANT-1 for treatment. **(E)** The BMS score at 4 weeks post crush injury (n =4 for Crush SCI, n = 3 for other three groups). **(F)** Immunostaining of Arl13b and Foxj1 showing the ependymal cells with ciliary features in the lesion site at 1 wpi, the details of the double-positive cells in the white squares are shown in the enlarged images (f1, f2, f3, f4, f5, f6, f7, f8). Scale bars, 50 μm on the top and 20 μm on the bottom. **(G)** Quantitative analysis of the proportion of Arl13b^+^Foxj1^+^ cells among Foxj1^+^ cells in the lesion site at 1 wpi for each group (n = 3 for Crush Noscapine+SANT-1, n = 4 for other three groups). **(H)** Immunostaining of GFAP and Foxj1 showing the ependymal cells exhibiting GFAP-labeled characteristics in the lesion site at 1 wpi, the details of the double-positive cells in the white squares are shown in the enlarged images (h1, h2, h3, h4, h5, h6, h7, h8). Scale bars, 50 μm on the top and 20 μm on the bottom. **(I)** Quantitative analysis of the proportion of GFAP^+^Foxj1^+^ cells among Foxj1^+^ cells in the lesion site at 1 wpi for each group (n = 3 per group). **(J)** Immunostaining of NeuN and PKD2L1 showing the survival neurons and CSF-cNs in the lesion site at 1 wpi, the details of CSF-cNs in the white squares are shown in the enlarged images (j1, j2, j3, j4). Scale bars, 200 μm on the top and 50 μm on the bottom. **(K)** Quantitative analysis of the proportion of PKD2L1^+^ cells among NeuN^+^ cells in the lesion site at 1 wpi for each group (n = 3 per group). i.p., intraperitoneal injection. Data are shown as mean ± SEM. ns, not significant (*P* > 0.05), \**P* < 0.05, \*\**P* < 0.01, \*\*\*\**P* < 0.0001. Unpaired two-sided Student’s t test.

Because Noscapine promotes ependymal differentiation lineage progression and enhances CSF-cNs survival-processes tightly associated with intact cilia-we examined whether these cellular effects were altered when cilia were disrupted. SANT-1 significantly suppressed the Noscapine-mediated increase in the proportion of GFAP⁺Foxj1⁺ ependymal cell-associated astrocytes (Figures 5H and 5I), indicating that ciliary function contributes to the lineage-associated transition of ependymal cells toward astrocytes. Likewise, the survival of CSF-cNs decreased under SANT-1 treatment, reflected by a reduced proportion of PKD2L1⁺ neurons (Figures 5J and 5K). These results suggest that Noscapine’s protective influence on CSF-cNs also depends on preserved ciliary integrity.

To further evaluate the role of cilia integrity in SCI repair, we turned to Y-27632, a Rho-associated protein kinase (ROCK) inhibitor reported to enhance ciliogenesis in vitro by altering actin dynamics^46^. Y-27632 has shown beneficial effects in SCI-including axon growth promotion and scar reduction-though its impact on cilia has not previously been examined^47–49^. In hTERT RPE-1 cells, Y-27632 increased cilia length and the proportion of ciliated cells (Figures 6A-6C), confirming its ciliogenesis-promoting activity in vitro. We next administered Y-27632 systemically after thoracic crush injury (Figure 6D). At 1 wpi, Y-27632 significantly increased the proportion of ciliated ependymal cells in vivo (Figures 6E and 6F), consistent with protection of ciliary structure during injury. Additionally, Y-27632 enhanced both the emergence of ependymal cell-associated astrocytes (Figures 6G and 6H) and CSF-cNs survival (Figures 6I and 6J), paralleling the effects of Noscapine and Paclitaxel. Together, these findings demonstrate that Noscapine’s reparative actions rely, at least in part, on preserved ciliary structure and function. Disrupting cilia-dependent signaling with SANT-1 attenuated Noscapine’s benefits, whereas enhancing cilia formation or stability with Y-27632 recapitulated them. These results support a mechanistic link between ciliary integrity, cellular responses, and functional recovery, highlighting cilia as a promising target for SCI intervention.

**Figure 6.**
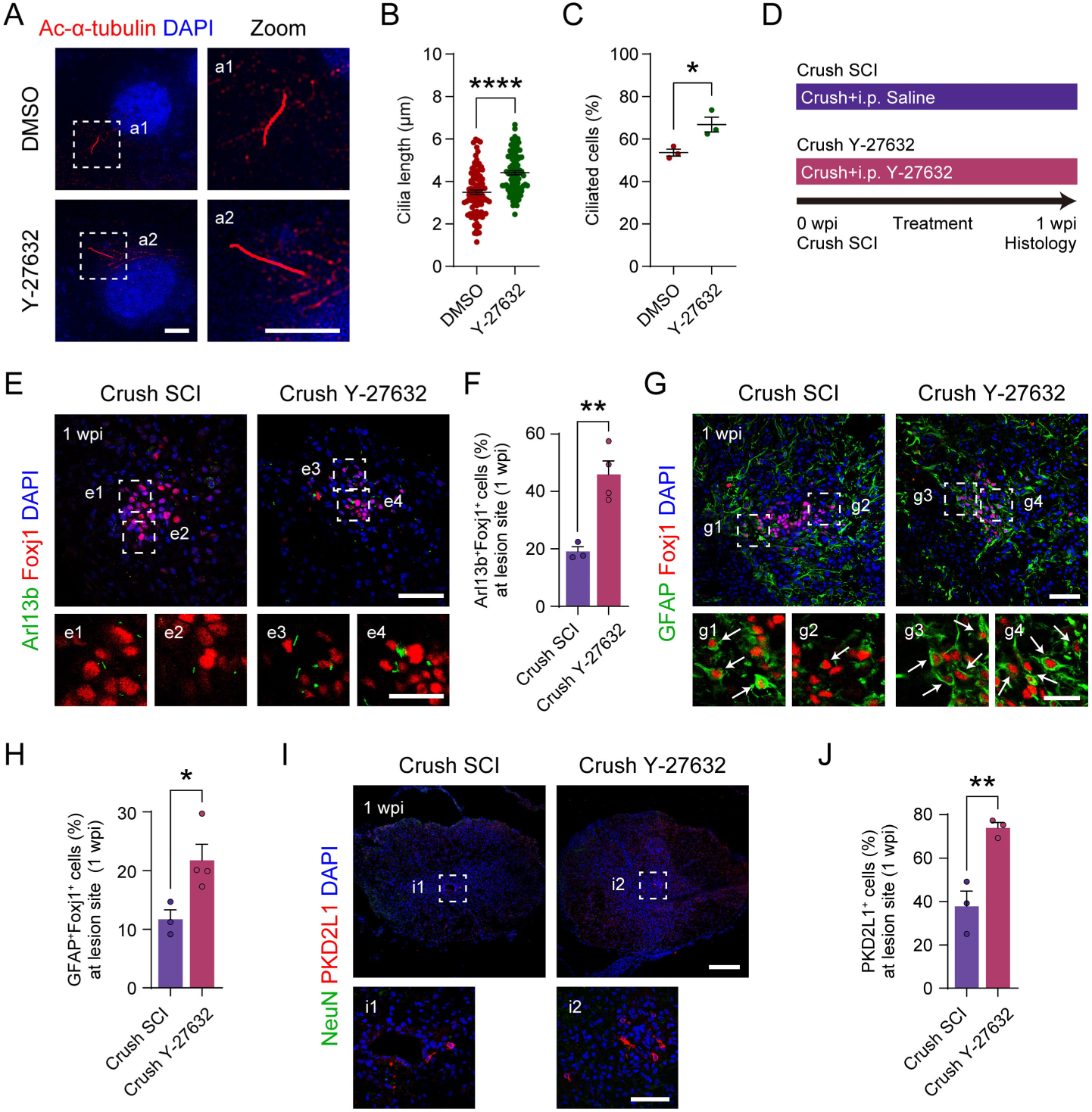
Ciliary integrity is a key factor in regulating the functional recovery from SCI. **(A)** Immunostaining of acetylated-tubulin (Ac-α-tubulin) showing the effect of Y-27632 on the cilia in hTERT-RPE1 cells. Scale bar, 5 μm. **(B)** Quantitative analysis of the length of cilia in each treatment (n = 100 per group). **(C)** Quantitative analysis of the percentage of ciliated cells in each treatment (n = 3 per group). **(D)** Diagram of experimental grouping and operation timeline using Y-27632 for treatment. **(E)** Immunostaining of Arl13b and Foxj1 showing the ependymal cells with ciliary features in the lesion site at 1 wpi, the details of the double-positive cells in the white squares are shown in the enlarged images (e1, e2, e3, e4). Scale bars, 50 μm on the top and 20 μm on the bottom. **(F)** Quantitative analysis of the proportion of Arl13b^+^Foxj1^+^ cells among Foxj1^+^ cells in the lesion site at 1 wpi for each group (n = 3 for Crush SCI, n = 4 for Crush Y-27632). **(G)** Immunostaining of GFAP and Foxj1 showing the ependymal cells exhibiting GFAP-labeled characteristics in the lesion site at 1 wpi, the details of the double-positive cells in the white squares are shown in the enlarged images (g1, g2, g3, g4). Scale bars, 50 μm on the top and 20 μm on the bottom. **(H)** Quantitative analysis of the proportion of GFAP^+^Foxj1^+^ cells among Foxj1^+^ cells in the lesion site at 1 wpi for each group (n = 3 for Crush SCI, n = 4 for Crush Y-27632). **(I)** Immunostaining of NeuN and PKD2L1 showing the survival neurons and CSF-cNs in the lesion site at 1 wpi, the details of CSF-cNs in the white squares are shown in the enlarged images (i1, i2). Scale bars, 200 μm on the top and 50 μm on the bottom. **(J)** Quantitative analysis of the proportion of PKD2L1^+^ cells among NeuN^+^ cells in the lesion site at 1 wpi for each group (n = 3 per group). i.p., intraperitoneal injection. Data are shown as mean ± SEM. \**P* < 0.05, \*\**P* < 0.01, \*\*\*\**P* < 0.0001. Unpaired two-sided Student’s t test.

## Discussion

Here, we identify preservation of ciliary integrity within central canal-associated cells as a structural mechanism that unifies the reparative actions of microtubule stabilizers after SCI. Despite acting through distinct chemotypes and tubulin-binding modes, Noscapine and Paclitaxel produced convergent structural and functional improvements across injury paradigms. Mechanistic analysis further revealed that ependymal cells and cerebrospinal fluid–contacting neurons represent principal responsive populations in which maintenance of ciliary architecture links microtubule stabilization to lineage-associated transition of ependymal cells and CSF-cNs survival. Importantly, attenuation of recovery under cilia-disrupting conditions establishes ciliary integrity as a shared structural checkpoint underlying cross-chemotype convergence and identifies a cilia-defined central canal regenerative niche as a mechanistic axis of spinal cord repair.

### Microtubule stabilization as a determinant of structural repair

Axonal degeneration and excessive fibrotic scarring jointly limit regeneration and circuit rebuilding after SCI, while microtubule destabilization represents a central barrier to structural repair^50^. Under a unified LOCS implantation and systemic dosing paradigm, both Noscapine and Paclitaxel reduced ECM/fibrotic scarring, increased neuronal presence, and improved axonal continuity, yielding convergent locomotor and electrophysiological benefits across injury paradigms (Figures 1 and 2). Brainbow-based anterograde circuit labeling with whole-cord clearing^20,51,52^, further revealed anatomical continuity across the lesion, although synaptic resolution remains limited (Figures 2 and S2)^53^.

Noscapine binds a colchicine-adjacent region rather than the taxane pocket^12^, yet displayed dose-dependent efficacy (Figure S1), reinforcing that stabilization of microtubule architecture-rather than chemotype-specific activity-drives structural repair. Given Paclitaxel’s cytotoxicity and poor solubility^13^, Noscapine’s favorable safety, solubility, and CNS penetration enhance the translational feasibility of a structure-centered stabilization strategy^54,55^. Importantly, extending beyond the classical view that microtubule stabilization primarily protects axonal structure^56^, our findings suggest that its reparative effects also involve preservation of a permissive central canal microenvironment that supports lineage-associated transition and neuronal survival after injury. Together, these observations establish microtubule stabilization as a chemotype-independent driver of structural repair and provide a mechanistic foundation for the cilia-dependent central canal regenerative niche identified in this study.

### Central canal-associated populations are key responders

Single-cell analysis revealed that, although Noscapine broadly modulates inflammatory and stromal compartments (Figures S4 and S5), its most coherent effects converged on two populations within the central canal niche. In ependymal cells, Noscapine enhanced progression toward astrocytes displaying neurotrophic and ensheathment-related signatures rather than scar-forming profiles (Figures 3 and S6). In parallel, Noscapine markedly improved the survival of CSF-cNs (Figure S7), a specialized sensory-modulatory population implicated in adhesion, angiogenesis, and injury-evoked tissue remodeling^42^.

These findings indicate that Noscapine does not merely reshape the lesion microenvironment but selectively engages the central canal niche, coordinating the emergence of ependymal cell-associated astrocytes and CSF-cNs preservation. The emergence of these two distinct canal-associated cell types as dominant responders-despite their differing developmental origins and functional roles—suggests that their responses may depend on a shared structural substrate within the central canal microenvironment that enables coordinated regeneration-associated remodeling after injury.

### Ciliary integrity links microtubule stabilization to lineage-associated cellular transition and neuronal support

GO enrichment analysis revealed that Noscapine upregulated cilia/IFT programs in both ependymal cells and CSF-cNs (Figures 4A, 4B, and 4E). Because SCI induces fragmentation of ependymal cilia^38,57,58^, the preservation of continuous ciliary organization in vivo (Figures 4C and 4D) indicates protection of these microtubule-based organelles. Noscapine similarly increased cilia-associated gene expression and preserved intact cilia in CSF-cNs (Figures 4E and 4F). In vitro, Noscapine enhanced cilia length and ciliation in RPE-1 cells (Figures 4G-4I), consistent with direct structural support outside the injury context. Paclitaxel exhibited similar trends in supporting ciliated ependymal cells and CSF-cNs survival (Figure S8), reinforcing ciliary integrity as a cross-chemotype structural effector of microtubule-based repair.

Together, these results indicate that ciliary integrity represents a downstream structural dependency through which microtubule stabilization supports lineage-associated cellular transition and neuronal preservation after injury.

### Ciliary preservation as a structure-centered potential repair strategy

Our findings position ciliary integrity as a critical structural determinant of SCI recovery and a unifying downstream dependency of microtubule stabilization. Pharmacological perturbation further supported the structural linkage: SANT-1 partially attenuated Noscapine’s effects on ependymal ciliation, lineage-associated transition, CSF-cNs survival, and motor recovery (Figure 5), whereas Y-27632 reproduced Noscapine-like improvements and increased ciliated ependymal cells in vivo (Figure 6).

Looking forward, cell type-specific and temporally controlled genetic approaches (e.g., Foxj1-CreER; Ift88/Kif3a) will be essential to test the necessity of cilia at cell differentiation^38,40^, and in vivo ciliary readouts (beat coordination, cerebrospinal fluid flow, IFT dynamics) will help link structural preservation to physiological function^37,59^. Because CSF-cNs integrate into local spinal circuits, future genetic tracing (e.g., Pkd2l1-Cre) may reveal whether their Noscapine-enhanced survival contributes directly to circuit-level reconstruction^41^.

Together, these perspectives support a cilia-centered framework in which microtubule stabilization promotes SCI recovery by maintaining ciliary architecture across central canal-associated cell types. Conceptually, this work shifts microtubule-targeted intervention from a compound-centered strategy toward a structure-centered paradigm; translationally, it elevates ciliary preservation from a correlative descriptor to a mechanistically grounded endpoint for guiding regeneration-oriented SCI interventions.

## Methods

### Animals

Adult female C57BL/6J mice (weight 18-20 g, age 6-8 weeks), purchased from Beijing Vital River Laboratory Animal Technology Co., Ltd. were used to establish the spinal cord injury model. Transgenic mouse Brainbow 3.2 (line 7) mice (Jackson Laboratory, stock #021227)^20^ were used for AAV tracing experiments. Use the following primers for PCR genotyping to detect WT or Brainbow 3.2 mice:

Transgene forward primer: CCACCTGATCTGCAACTTGA

Transgene reverse primer: TGCTAGGGAGGTCGCAGTAT

Wild type forward primer: CTAGGCCACAGAATTGAAAGATCT

Wild type reverse primer: GTAGGTGGAAATTCTAGCATCATCC

All animals were housed under specific pathogen-free (SPF) conditions at the Laboratory Animal Center, Institute of Genetics and Developmental Biology, Chinese Academy of Sciences. The animals were given water and food for 24 hours under standard housing conditions (12-h light/12-h dark cycle). All animal experimental procedures were performed in accordance with the guidelines of the Animal Ethics Committee of the Institute of Genetics and Developmental Biology, Chinese Academy of Sciences.

### Antibodies and Reagents

The following primary antibodies were used: mouse anti-EB1 (BD Transduction Laboratories, 610535; 1:250), chicken anti-GFAP (Abcam, ab4674; 1:1000), rabbit anti-Laminin (Sigma-Aldrich, L9393; 1:100), mouse anti-NeuN (Millipore, MAB377; 1:500), mouse anti-SMI-312 (Biolegend, 837904; 1:500), chicken anti-GFP (Invitrogen, A10262; 1:500), rabbit anti-tRFP (Evrogen, AB233, 1:500), guinea pig anti-mCherry (Oasis Biofarm, OB-PGP004; 1:500), rabbit anti-PKD2L1 (Millipore, AB9084; 1:500), mouse anti-Foxj1 (Invitrogen, 14-9965-82; 1:200), rabbit anti-Arl13b (Proteintech, 17711-1-AP; 1:500), mouse anti-Ac-α-tubulin (Sigma-Aldrich, T6793; 1:500).

The following Secondary antibodies were used: Goat anti-Mouse, Alexa Fluor 488, 555, 647 (Thermo Fisher Scientific); Goat anti-Rabbit, Alexa Fluor 555, 647 (Thermo Fisher Scientific); Goat anti-Guinea Pig, Alexa Fluor 647 (Invitrogen); Donkey anti-Chicken, Alexa Fluor 488 (Jackson).

Reagents: Noscapine hydrochloride hydrate (Macklin, 912-60-7); Epothilone D (APExBIO, 189453-10-9); TPI-287 (Bioruler, 849213-15-6); TTI-237 (AdooQ, 849550-05-6); Paclitaxel (MedChemExpress, 33069-62-4); Paclitaxel liposome for injection (Nanjing Luye Pharmaceutical Co., Ltd); SANT-1 (Aladdin, 304909-07-7); Y-27632 dihydrochloride (MedChemExpress, 129830-38-2); DMSO (Sigma-Aldrich, 67-68-5).

### Cell culture and drug treatments

HeLa cells (obtained from Masatoshi Takeichi, RIKEN Center for Biosystems Dynamics Research, Japan; RRID: CVCL_0030) and hTERT-RPE1 cells (ATCC; RRID: CVCL_4388) were cultured in DMEM/F12 1:1 (Wisent) medium supplemented with 10% fetal bovine serum (FBS, TransSerum) and 1% penicillin/streptomycin (P/S, Wisent) at 37℃, 5% CO_2_. HeLa cells were treated with the following compounds of DMSO, 50 μM Noscapine, 10 μM Epothilone D, 10 μM TPI-287, 10 μM TTI-237, 10 μM Paclitaxel for 2 h to assess microtubule dynamics.

hTERT-RPE1 cells were grown to approximately 90% confluency. Cells were then washed twice with phosphate-buffered saline (PBS) and incubated in DMEM/F12 (1:1) medium without FBS for 24-48 hours to induce ciliogenesis. The ciliated hTERT-RPE1 cells were subsequently treated with either DMSO, 50 μM Noscapine, 10 μM SANT-1 or 40 μM Y-27632 for 24 hours.

### Immunocytochemistry

Following drug treatment, cells were washed twice with PBS and fixed with 4% paraformaldehyde (PFA) at 37°C for 10 min. Cells were then washed three times for 5 min each with PBST (0.1% Triton X-100 in PBS). Subsequently, cells were blocked with 3% bovine serum albumin (BSA) at room temperature for 30 min. After blocking, cells were incubated with primary antibodies at room temperature for 1 hour, followed by three 5-min washes with PBST. Cells were then immediately incubated with appropriate secondary antibodies at room temperature for 1 hour, followed by three 5-min washes with PBS. Finally, the cells were mounted on slides using FluorSave™ reagent (Millipore). When performing immunostaining with the anti-EB1 antibody, cells were fixed with absolute methanol at -20°C for 5 min prior to the subsequent steps. Images of stained cells were photographed using the Multi-SIM imaging system.

### Spinal cord injury surgery

Female adult mice were deeply anesthetized by intraperitoneal injection of Avertin (Sigma-Aldrich, 75-80-9) at a dose of 20 μL/g. Mice were secured in a prone position, the dorsal fur was shaved, and the skin was disinfected with iodine solution. A midline incision was made above the thoracic vertebrae with a scalpel. Muscle tissue was dissected to expose the T8-T9 vertebral segments, and a laminectomy was performed. For thoracic complete transection spinal cord injury, the 1-2 mm exposed spinal cord segment was transected and removed using micro-scissors. The linear-ordered collagen scaffold (LOCS), loaded with drug, was then transplanted into the lesion epicenter. For thoracic crush spinal cord injury, carefully insert the forceps into both sides of the exposed spinal cord and completely clamp the spinal cord for 10 seconds. After injury, the muscle layer and skin were sutured sequentially. Mice were placed on a heating pad until full recovery from anesthesia. Penicillin (SANTAIBIO, GC-qms-1) was injected for 5 consecutive days, and artificial urination was maintained twice a day until tissue collection.

During the experiment, in the thoracic complete transection spinal cord injury, the LOCS was soaked in either saline or Noscapine for 10 min and then transplanted to the lesion site. Mice subsequently received daily intraperitoneal injections of either saline or Noscapine until tissue collection. In the thoracic crush spinal cord injury, mice received daily intraperitoneal injections of either saline, Noscapine, Paclitaxel or SANT-1 until tissue collection.

### Behavioral assessment

The recovery of hindleg locomotor function was assessed weekly from 1 to 8 weeks post-injury using the Basso Mouse Scale (BMS) score^60^ by two independent observers blinded to the experimental groups.

During weekly locomotor assessments, mice activity in the open field was recorded using a camera system. Recordings were analyzed in Adobe Premiere Pro 2021 to observe hindleg movement throughout the time series. Key frames before and after the mice’s hindleg swing were selected and exported as images. By drawing the tangent of the hindleg calf muscle and the direction of the claws in each key frame, the angle between the two was measured, and calculate the difference in this angle between pre-swing and post-swing frames to represent hindleg movement extent.

### Electrophysiology assessment

Cortical motor evoked potentials (CMEPs) were recorded using Keypoint 9033A07 (Alpine bioMed ApS, Denmark) at 4 and 8 weeks post-injury. Mice were deeply anesthetized with Avertin. A stimulating electrode was placed into the subcutaneous tissue 1 cm behind the intersection of the skull midline and the eyebrows. A recording electrode was inserted into the muscle of one hindleg and a ground electrode was placed in the ventral subcutaneous tissue. The latency and amplitude, obtained following multiple-pulse stimulation, were measured to evaluate the recovery level of the hindleg.

### Tissue collecting and immunostaining

At 1 or 4 weeks post-injury, deeply anesthetized mice underwent transcardial perfusion with PBS followed by 4% PFA. Approximately 2 cm of spinal cord tissue was removed from the center of the injury. The tissue was then post-fixed in 4% PFA at 4°C overnight.

Subsequently, the tissue was submerged in 30% sucrose at 4°C for 24 h. The tissue was then embedded in OCT (Sakura, 4583) and stored at -80°C until use. Cryosections of 10-20 μm were prepared in longitudinal or transverse of spinal cord tissue using LEICA CM 1950 cryostat and stored at -20°C.

Cryosections were washed three times for 5 min each with PBS at room temperature, and then blocked with 5% BSA in PBS for 30 min. Following blocking, sections were incubated with primary antibodies overnight at 4°C. After washing with PBS, the sections were incubated with appropriate secondary antibodies for 1 hour at room temperature. After three additional 5-min washes with PBS, sections were coverslipped using mounting medium. For immunostaining with anti-DCX and anti-Foxj1 antibodies, antigen retrieval was performed using citrate repair solution (ORIGENE, ZLI-9064) for 30 min in a 95°C water bath prior to blocking. Fluorescence images were acquired using an Olympus FV3000RS confocal microscope.

### Anterograde tracing in Brainbow mice

Mice were deeply anesthetized and underwent a laminectomy at T6-T9. AAV2/1-hSyn-Cre (1×10^13^ V.G./mL, Shanghai Taitool Bioscience Co., Ltd) was bilaterally (0.3 mm) injected into the spinal cord at T6. Injections were made at two depths (0.9 mm and 0.4 mm below the dorsal surface). A total of 100 nL was injected per site at a rate of 50 nL/min. Following viral injection, a complete transection spinal cord injury was performed at T8-T9, and mice were perfused for tissue collection 4 weeks post-injury. The fluorescent proteins EGFP, mKate2, and mOrange2 in the spinal cord were labeled by immunostaining with chicken anti-GFP, rabbit anti-tRFP, and guinea pig anti-mCherry.

### Tissue Clearing and photograph

Four weeks post-viral injection, mice were perfused transcardially, and the injured site tissue was collected, fixed overnight at 4°C with 4% PFA, followed by carefully remove the spinal dura mater. To enhance fluorescent labeling, whole-mount immunostaining of the intact spinal cord tissue was performed, adapted from the method described by Hilton et al.^52^. Tissue was incubated in PBS containing 0.2% Triton X-100, 20% DMSO, and 0.3 M glycine at 37°C with shaking overnight. Then, tissue was blocked in PBS containing 0.2% Triton X-100, 10% DMSO, and 6% BSA overnight at 37°C. Tissue was washed twice for 1 hour each with PBS containing 0.2% Tween 20 and 10 U/ml heparin (PTwH). The primary antibodies were incubated with shaking at 37°C for three days. Tissue was then washed twice with PTwH for 6 hours each, followed by an additional overnight wash in fresh PTwH. The appropriate secondary antibodies were shaken at 37°C for two days. Finally, tissue was washed extensively with PTwH for 2 days (with solution changes every 12 hours). The clearing procedure was performed as previously described^61^. Briefly, tissue was incubated overnight in 50% tetrahydrofuran (THF, Macklin, 109-99-9), followed by incubation in 80% THF for 1 hour. After two 1-hour incubations in 100% THF, tissue was transferred to 100% dichloromethane (Macklin, 75-09-2) until it sank to the bottom. Finally, tissue was submerged in BABB clearing solution (a 1:2 mixture of benzyl alcohol (Macklin, 100-51-6) and benzyl benzoate (Macklin, 120-51-4)) until optically transparent, and then mounted on slides containing BABB for microscopic imaging. All the clearing processes were conducted at room temperature in the dark. Samples were imaged using a two-photon microscope (Two-photon Nikon A1R MP system). Image processing and 3D reconstruction were performed using IMARIS 9 software (Bitplane, AG).

### Single-cell suspension preparation

Following deep anesthesia, mice were perfused transcardially with saline. Collecting the injury site tissue and carefully removing the spinal dura mater in cold PBS. Tissue dissociation was performed using the Neural Tissue Dissociation Kit (Miltenyi Biotec, 130-092-628) according to the manufacturer’s protocol. Briefly, the tissue was cut into small pieces and digested in tissue dissociation solution at 37°C for 1.5 hours. The digestion was stopped by adding an equal volume of DMEM/F12. The cell suspension was sequentially filtered through 100 μm and 40 μm cell strainers, and cell pellet was collected by centrifugation at 400 × *g* for 5 min. The cell pellet was resuspended in 25% Percoll (Solarbio, P8370) and centrifuged at 400 × *g* for 10 min to remove myelin debris. The cell pellet was then resuspended in Red Blood Cell Lysis Buffer (Solarbio, R1010). After standing 2 min at room temperature, an equal volume of saline was added to stop lysis. Cells were centrifuged at 400 × *g* for 5 min and the pellet was resuspended in saline. Propidium Iodide (PI) (Sigma-Aldrich, 25535-16-4) was added to the cell suspension, and dead cells and cell debris were removed using a flow cytometer (BD FACSAria II). Sorted cells were collected in PBS containing 5% BSA, centrifuged at 2000 × *g* for 5 min, and finally resuspended in saline. The cell number and viability in the cell suspension were checked under a microscope.

### Single-cell RNA sequencing

Single-cell cDNA library construction and sequencing were performed according to the official library construction process and standards of 10X Genomics. In the Chromium™ microfluidic system, each cell was encased in separated oil droplets with gel beads containing barcodes to form Gel Beads-In-Emulsion (GEMs). In each GEM, mRNA released by cell lysis was reverse-transcribed into cDNA with barcodes. Then the oil droplets were broken, and the barcoded cDNA was collected to construct a next-generation sequencing library. The library quality was assessed using an Agilent 2100. Finally, the Illumina NovaSeq 6000 platform was used to sequence the raw sequencing data for downstream analysis. To ensure the reliability and accuracy of the analysis results, the multiple cells predicted by Scrublet software were removed, and low-quality cells with less than 200 genes or the total expression of all genes less than 400 or the proportion of mitochondrial genes was more than 10% were filtered out.

Clustering and differential gene expression analysis were performed using Seurat. The FindVariableFeatures function was used to identify the top 2000 highly variable genes (HVGs) among cells. The FindIntegrationAnchors and IntegrateData functions were applied with a global normalization method to eliminate batch effects between samples. Dimensionality reduction and clustering were then performed using the ScaleData function and the results were visualized using Uniform Manifold Approximation and Projection (UMAP).

Subsequent analysis was performed using Seurat and our custom R scripts. The FindAllMarkers function was used for differential gene expression analysis. All cell types in the samples were manually annotated based on the expression level of classical marker genes combined with the differential expression results. To perform further sub-clustering analysis on specific cell type, the targeted cell types were re-clustered using Seurat, and the differential gene expression analysis (as described above) was repeated. Gene Ontology (GO) enrichment analysis was performed on the differentially expressed genes (DEGs) for each cell type across all samples using the ClusterProfiler package. Pseudo-time analysis of lineage differentiation between cell types of interest was conducted using monocle version 2.3.6, generating an intuitive lineage trajectory diagram.

### Quantification and statistical analysis

Fluorescence images from immunostaining were analyzed using ImageJ software to measure the mean fluorescent area and count positive cells. Statistical analysis and graphical generation were performed using GraphPad Prism 8.0, and all data were represented by mean ± SEM. The two-tailed Student’s t-tests were used to analyze the differences between two groups. The significance levels are denoted as follows: ns, not significant (*P* > 0.05), \**P* < 0.05, \*\**P* < 0.01, \*\*\**P* < 0.001, and \*\*\*\**P* < 0.0001. All experiments, as described in the figure legends, were performed with at least three biological replicates.

## Data availability

Single-cell RNA sequencing data in this study have been deposited to the ArrayExpress database under the accession number E-MTAB-15813. The published sequencing data is downloaded from Gene Expression Omnibus (GSE198949).

## Acknowledgements

We thank Zhengrong Zhou and Jinhui Shao for advice; Yongheng Fan for assistance with scRNA-seq experiments; Zhiheng Xu and Yisheng Jiang for assistance with microinjection instrument; Kangmin He for gifting hTERT-RPE1 cells. We thank Yan Teng, Qing Bian, Duo Duo, Xing Jia and Yun Feng for technical support with Confocal imaging.

## Author contributions

C.L. and W.M. designed all experiments; C.L. and X.W. analyzed the scRNA-seq data; C.L., L.Y., R.Z. and R.L. carried out the spinal cord injury modeling; C.L. and L.Y. performed the scRNA-seq experiments; C.L., R.Z. and H.X. analyzed image data; Q.X., Y.Z. and J.D. provided reagents and advice; C.L. and W.M. prepared the manuscript.

