## Supplemental information for "Ciliary integrity defines a central canal repair checkpoint linking microtubule stabilization to spinal cord regeneration"

**Supplementary information**

**Figure S1. Intraperitoneal injection of 100 mg/kg of noscapine produced a significant BMS improvement at 4 wpi.**

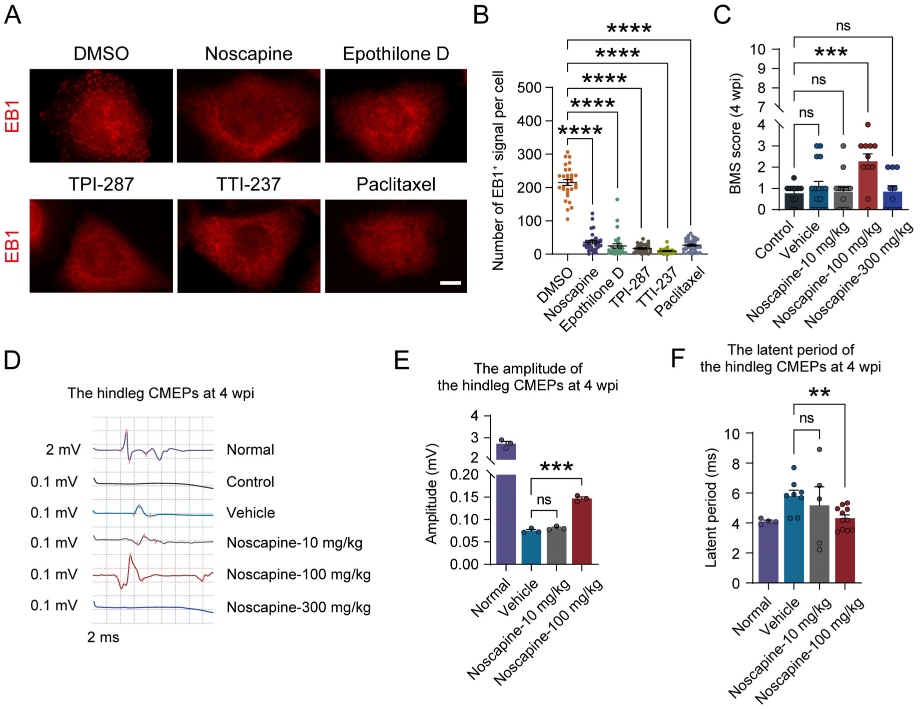


**(A)** Immunostaining for EB1 showing the effects of all compounds on the plus-end of microtubules in HeLa cells. Scale bar, 10 μm. **(B)** Quantitative analysis of the number of $EB1^+$ signal in each cell (n = 30 per group). **(C)** The BMS score at 4 wpi with different intraperitoneal injection dose (n = 11 for Control, n = 16 for Vehicle, n = 16 for Noscapine-10 mg/kg, n = 11 for Noscapine-100 mg/kg, n = 10 for Noscapine-300 mg/kg). **(D)** The representative images of cortical motor evoked potentials (CMEPs) between normal mice and experimental group mice at 4 wpi. **(E-F)** Quantitative analysis of the amplitude **(E)** (n = 3 per group) and latent period **(F)** (n = 4 for Normal, n = 8 for Vehicle, n = 5 for Noscapine-10 mg/kg, n = 11 for Noscapine-100 mg/kg) of the hindleg CMEPs at 4 wpi. CMEPs were not detected in the Control and Noscapine-300 mg/kg groups at 4 wpi, so they were not shown in figures. wpi, weeks post-injury. Data are shown as mean ± SEM. ns, not significant ($P > 0.05$), $^{**}P < 0.01$, $^{***}P < 0.001$, $^{****}P < 0.0001$. Unpaired two-sided Student's t test.

**Figure S2. Brianbow mice combined with AAV virus tracing strategy.**

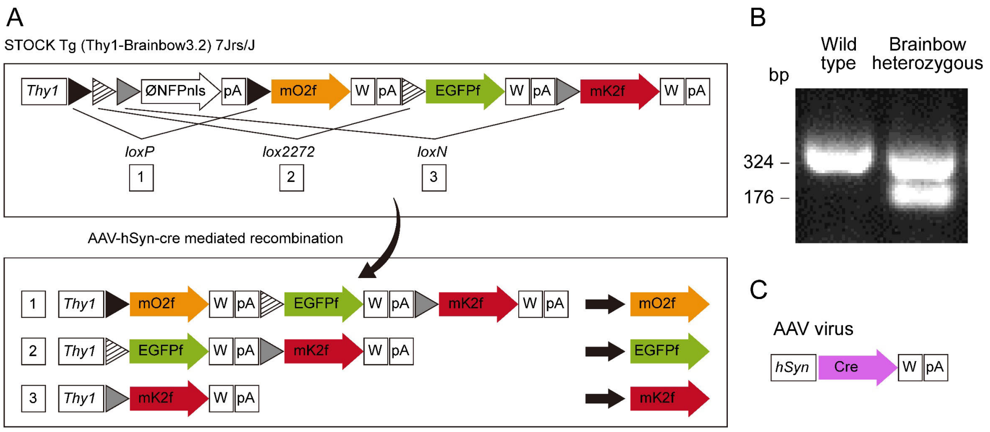


**(A)** The gene construct of Brainbow 3.2 transgenic mice. After Cre recombination, EGFP, mKate2, or mOrange2 will be expressed in an unpredictable manner, resulting in multiple fluorescent proteins being expressed within a single cell, producing various colors. **(B)** Genotyping of Brainbow 3.2 transgenic mice by PCR of genomic DNA. **(C)** The gene construct of AAV virus specifically expressing Cre in neurons. ØNFPnls, nuclear-targeted nonfluorescent XFP, mO2f, mOrange2f, mK2f, mKate2f, W, woodchuck hepatitis virus posttranscriptional regulatory element, pA, polyadenylation sequence.

**Figure S3. Single-cell RNA sequencing sample preparation and quality control.**

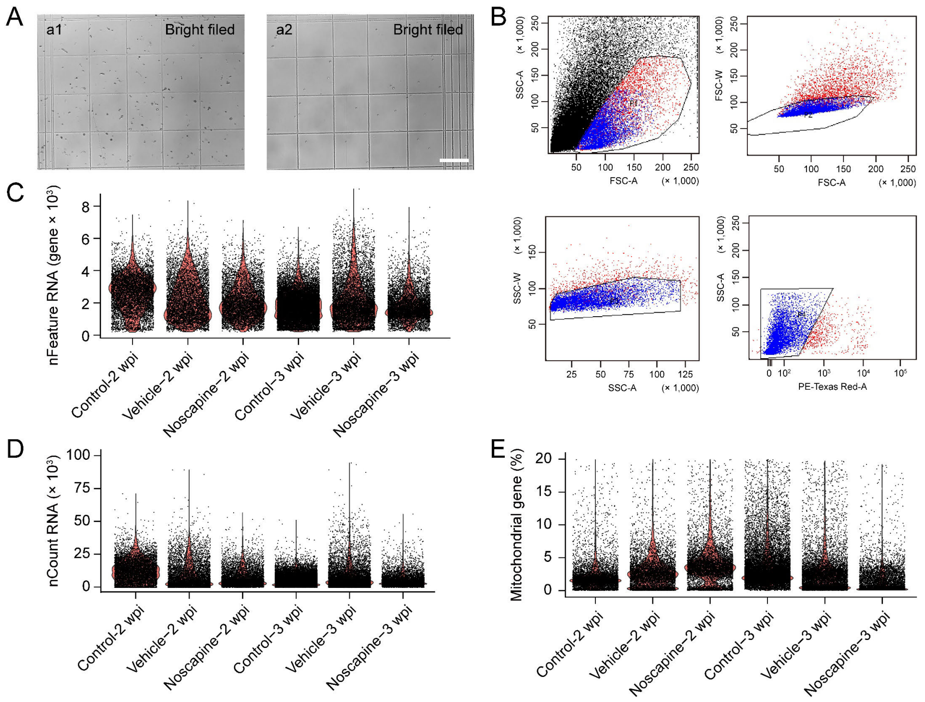


**(A)** Representative images of the cell suspension before (a1) and after (a2) fluorescence-activated cell sorting (FACS). Scale bar, 200 μm. **(B)** Dead cells and cellular debris were removed by FACS. Blue cells were selected live cells. **(C-E)** Retained high quality cells in each of the following metrics: cells with more than 200 genes **(C)**, cells with more than 400 UMI counts **(D)** and cells where the proportion of mitochondrial genes is less than 20% **(E)**. FSC, forward-scattered light, SSC, side-scattered light, -W, width, -A, area.

**Figure S4. Single-cell RNA sequencing analysis of the spinal cord lesion site tissue at 2 wpi and 3 wpi.**

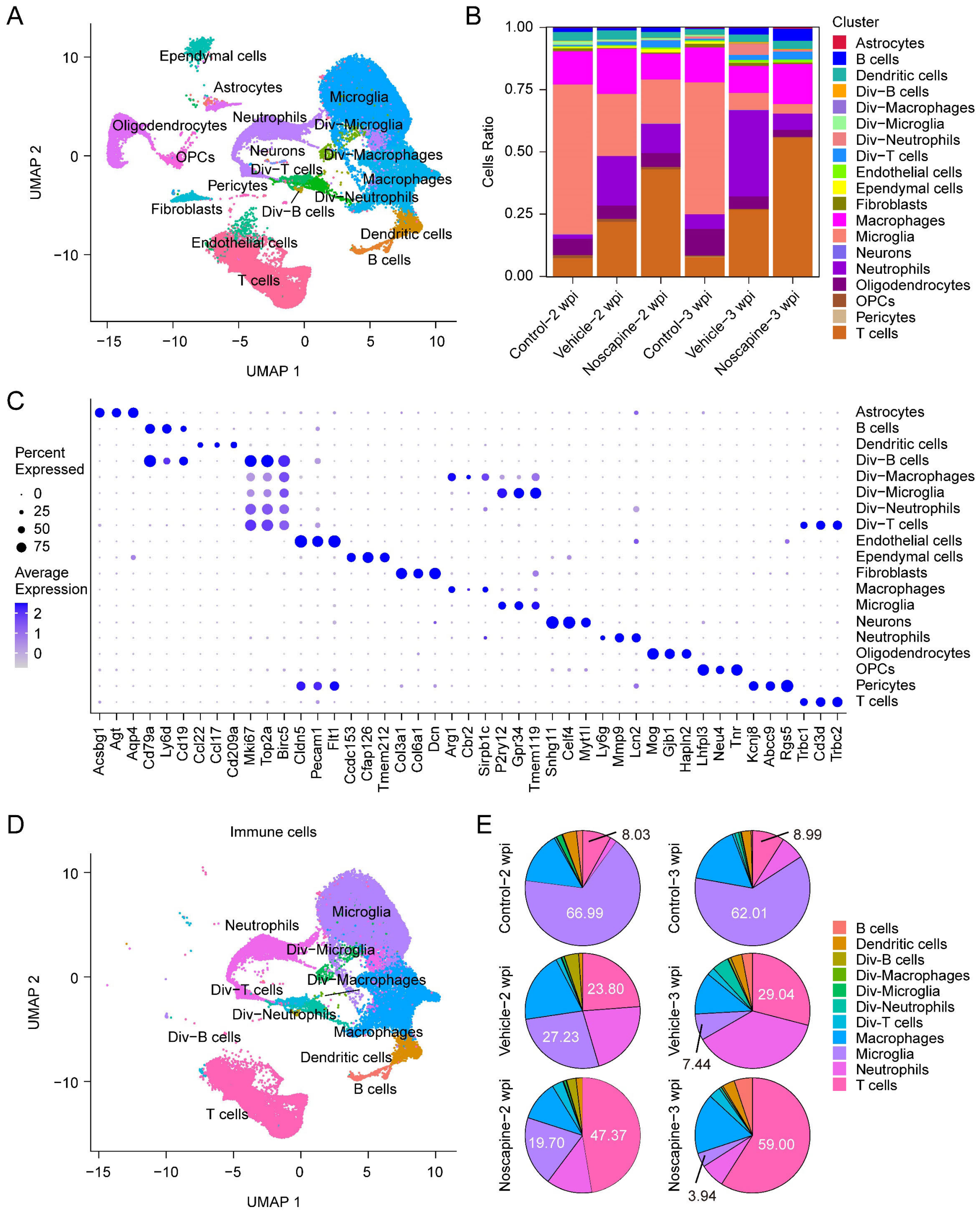


**(A)** UMAP plot showing 51,117 high-quality cell transcriptome profiles of six groups. **(B)** The dynamic changes in the percentage of cells in six groups. **(C)** Dot plot showing marker genes of cells in the spinal cord. The dot size indicates the percentage of cells expressing the gene in each cluster, and the color represents the average expression level. **(D)** UMAP plot showing immune cells of six groups. **(E)** Pie charts showing the percentage of each immune cell type across different groups. Div-, dividing-.

**Figure S5. Noscapine reduces the abundance of fibroblasts associated with extracellular matrix formation at lesion site.**

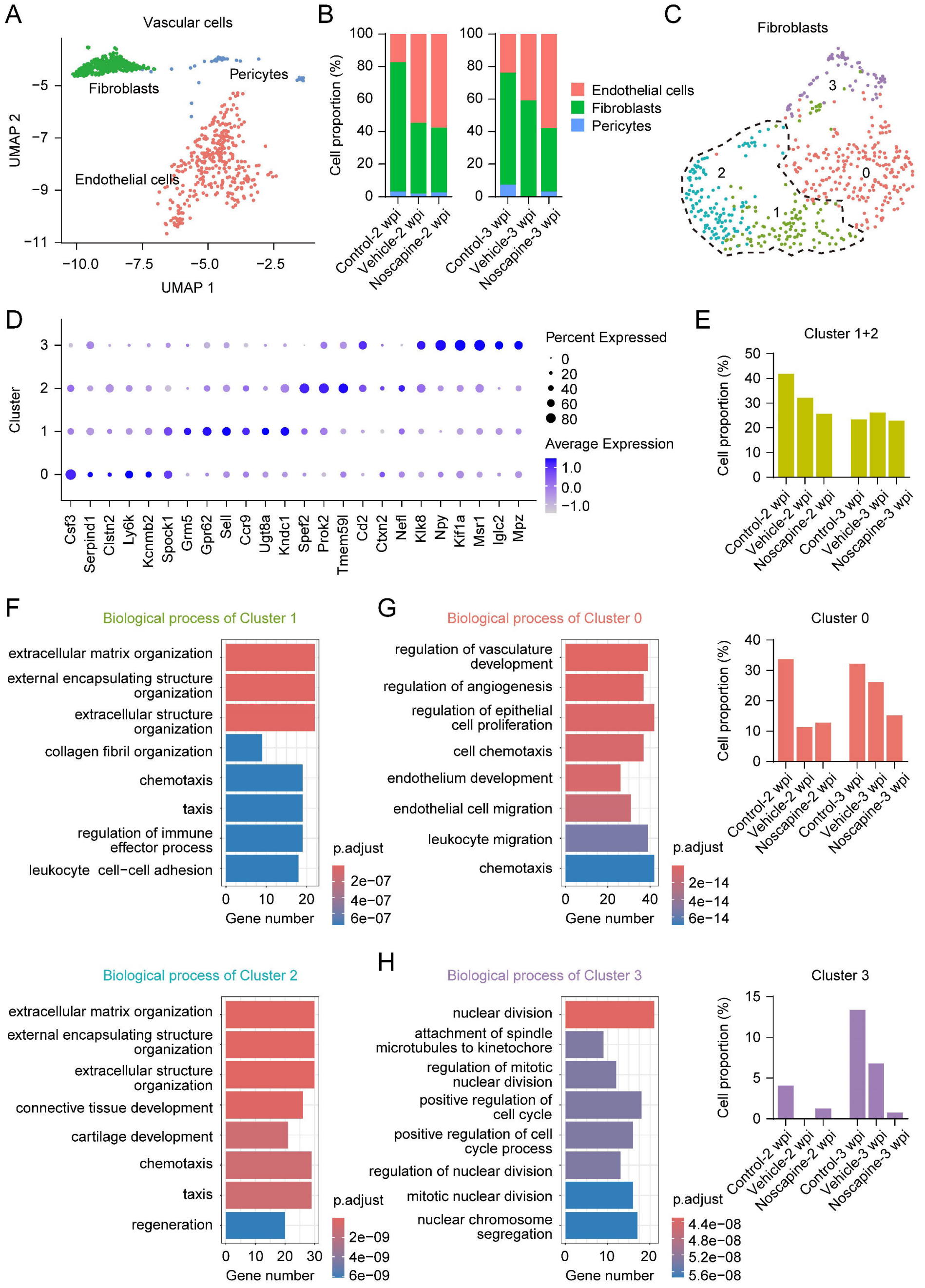


**(A)** UMAP plot showing vascular cells (fibroblasts, pericytes, endothelial cells) of six groups. **(B)** The percentage of vascular cells in each group. **(C)** UMAP plot showing fibroblast subpopulations of six groups. **(D)** Dot plot showing differentially expressed genes in each subpopulation. The dot size indicates the percentage of cells expressing the gene in each cluster, and the color represents the average expression level. **(E)** The percentage of cluster 1 and cluster 2 cells in each group. **(F)** GO enrichment analysis of biological process for the highly expressed genes in cluster 1 and 2. **(G)** GO enrichment analysis of biological process for the highly expressed genes in cluster 0 and cell proportion in each group. **(H)** GO enrichment analysis of biological process for the highly expressed genes in cluster 3 and cell proportion in each group.

**Figure S6. Cell composition of neurocyte cells pseudotime analysis and crush model establishment.**

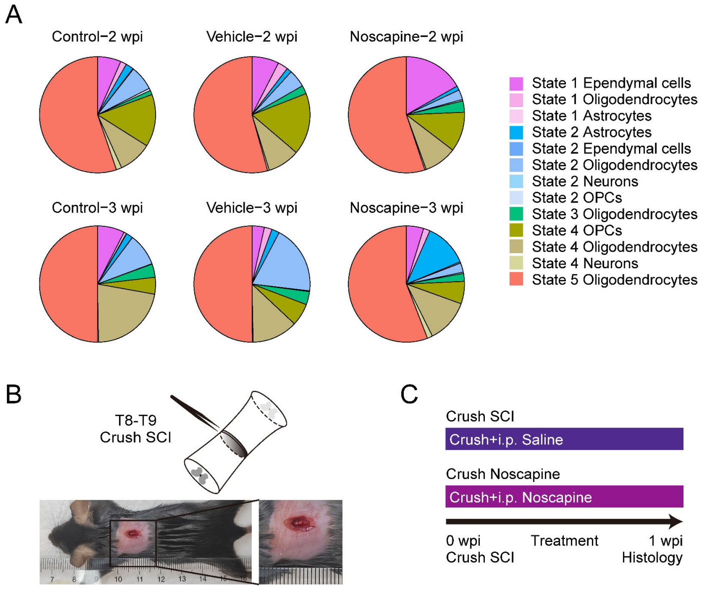


**(A)** Cell composition of neurocyte cells pseudotime analysis of six groups. **(B)** The diagram of surgical procedures of thoracic crush spinal cord injury. **(C)** Diagram of experimental grouping and operation timeline using noscapine for treatment. i.p., intraperitoneal injection.

**Figure S7. Noscapine treatment improves the survival of CSF-cNs.**

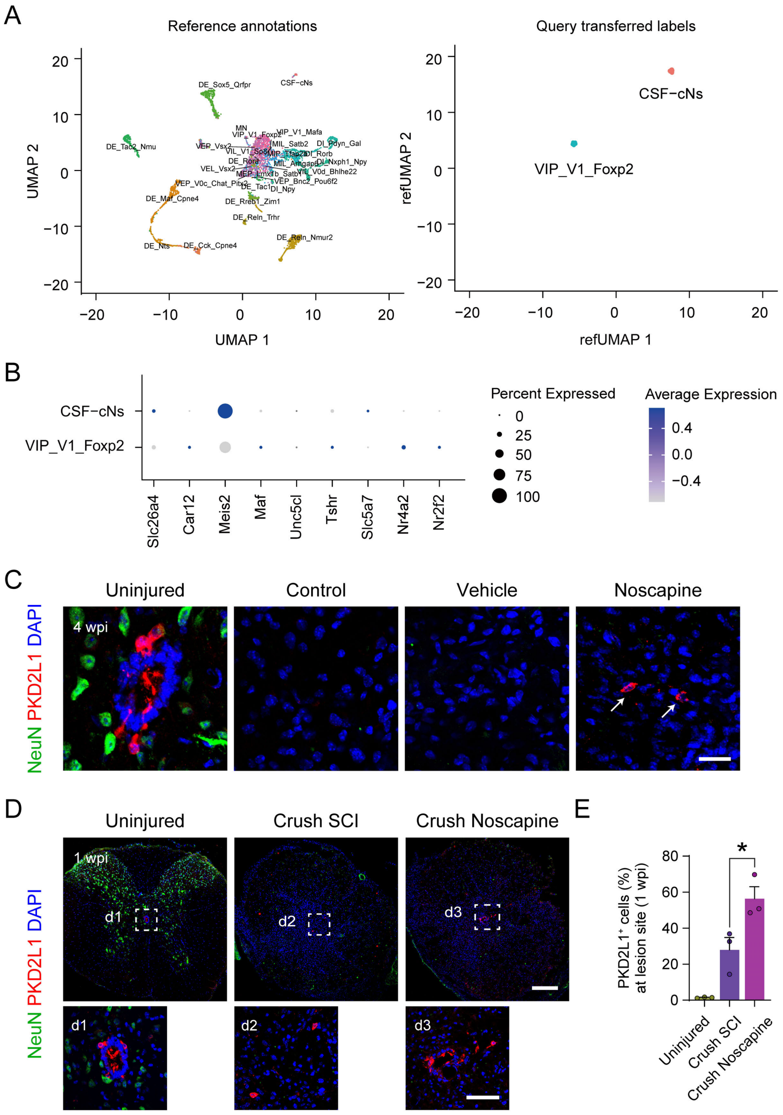


**(A)** UMAP plot showing neuron subsets identified using the thoracic spinal neuron atlas. **(B)** Dot plot showing marker genes of neurons in the spinal cord. The dot size indicates the percentage of cells expressing the gene in each cluster, and the color represents the average expression level. **(C)** Immunostaining of NeuN and PKD2L1 showing the survival neurons and CSF-cNs in the lesion site at 4 weeks post complete transection injury. Scale bar, 20 μm. **(D)** Immunostaining of NeuN and PKD2L1 showing the survival neurons and CSF-cNs in the lesion site at 1 wpi, the details of CSF-cNs in the white squares are shown in the enlarged images (d1, d2, d3). Scale bars, 200 μm on the top and 50 μm on the bottom. **(E)** Quantitative analysis of the proportion of $PKD2L1^+$ cells among $NeuN^+$ cells in the lesion site at 1 wpi for each group (n = 3 per group). Data are shown as mean ± SEM. $^*P < 0.05$. Unpaired two-sided Student's t test.

**Figure S8. Paclitaxel recapitulated cilia-linked effects in the crush model.**

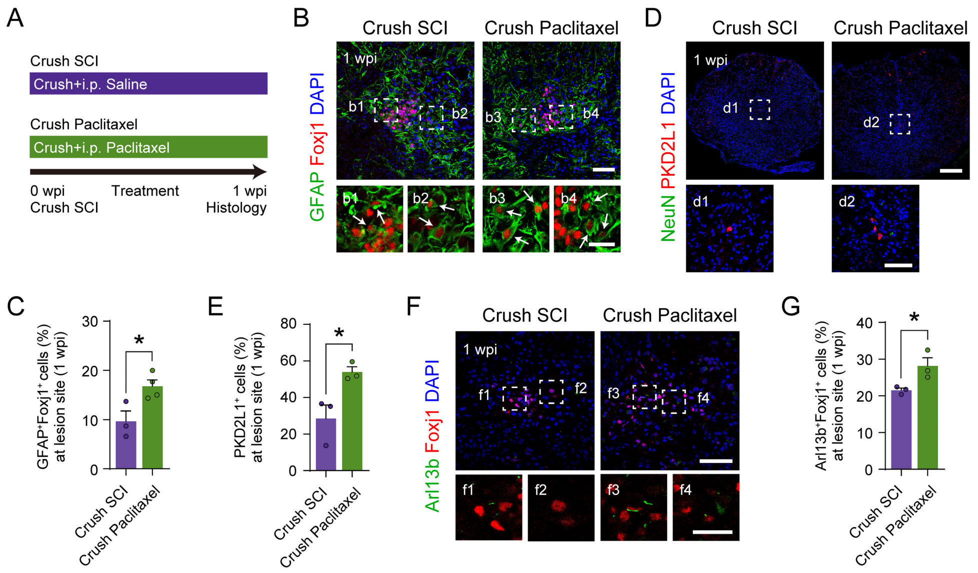


**(A)** Diagram of experimental grouping and operation timeline using paclitaxel for treatment. **(B)** Immunostaining of GFAP and Foxj1 showing the ependymal cells exhibiting GFAP-labeled characteristics in the lesion site at 1 wpi, the details of the double-positive cells in the white squares are shown in the enlarged images (b1, b2, b3, b4). Scale bars, 50 μm on the top and 20 μm on the bottom. **(C)** Quantitative analysis of the proportion of $GFAP^+Foxj1^+$ cells among $Foxj1^+$ cells in the lesion site at 1 wpi for each group (n = 3 for Crush SCI, n = 4 for Crush Paclitaxel). **(D)** Immunostaining of NeuN and PKD2L1 showing the survival neurons and CSF-cNs in the lesion site at 1 wpi, the details of CSF-cNs in the white squares are shown in the enlarged images (d1, d2). Scale bars, 200 μm on the top and 50 μm on the bottom. **(E)** Quantitative analysis of the proportion of $PKD2L1^+$ cells among $NeuN^+$ cells in the lesion site at 1 wpi for each group (n = 3 per group). **(F)** Immunostaining of Arl13b and Foxj1 showing the ependymal cells with ciliary features in the lesion site at 1 wpi, the details of the double-positive cells in the white squares are shown in the enlarged images (f1, f2, f3, f4). Scale bars, 50 μm on the top and 20 μm on the bottom. **(G)** Quantitative analysis of the proportion of $Arl13b^+Foxj1^+$ cells among $Foxj1^+$ cells in the lesion site at 1 wpi for each group (n = 3 per group). i.p., intraperitoneal injection. Data are shown as mean ± SEM. *$P < 0.05$. Unpaired two-sided Student's t test.
